# The natural frequencies of the resting human brain: an MEG-based atlas

**DOI:** 10.1101/2021.11.17.468973

**Authors:** Almudena Capilla, Lydia Arana, Marta García-Huéscar, María Melcón, Joachim Gross, Pablo Campo

## Abstract

Brain oscillations are considered to play a pivotal role in neural communication. However, detailed information regarding the typical oscillatory patterns of individual brain regions is surprisingly scarce. In this study we applied a multivariate data-driven approach to create an atlas of the natural frequencies of the resting human brain on a voxel-by-voxel basis. We analysed resting-state magnetoencephalography (MEG) data from 128 healthy adult volunteers obtained from the Open MEG Archive (OMEGA). Spectral power was computed in source space in 500 ms steps for 82 frequency bins logarithmically spaced from 1.7 to 99.5 Hz. We then applied k-means clustering to detect characteristic spectral profiles and to eventually identify the natural frequency of each voxel. Our results revealed a region-specific organisation of intrinsic oscillatory activity, following both a medial-to-lateral and a posterior-to-anterior gradient of increasing frequency. In particular, medial fronto-temporal regions were characterised by slow rhythms (delta/theta). Posterior regions presented natural frequencies in the alpha band, although with differentiated generators in the precuneus and in sensory-specific cortices (i.e., visual and auditory). Somatomotor regions were distinguished by the mu rhythm, while the lateral prefrontal cortex was characterised by oscillations in the high beta range (>20 Hz). Importantly, the brain map of natural frequencies was highly replicable in two independent subsamples of individuals. To the best of our knowledge, this is the most comprehensive atlas of ongoing oscillatory activity performed to date. Furthermore, the identification of natural frequencies is a fundamental step towards a better understanding of the functional architecture of the human brain.

## Introduction

Brain oscillations are spontaneously generated at rest and can be captured by means of electro- and magnetoencephalography (EEG/MEG) (for a review see Lopes da Silva, 2013). Critically, the fluctuations in electromagnetic brain activity are considered to be the main communication code between neural networks (Buzsáki & Watson, 2012; Fries, 2015; Varela et al., 2001). Hence, characterising the typical patterns of oscillatory activity is crucial for understanding the functional architecture of the brain that is at the core of perception, cognition, and behaviour.

Since Jasper and Penfield (1949) first attempts, several studies have tried to establish common oscillatory patterns across the resting human brain. Overall, oscillations at alpha frequency (8–13 Hz) are consistently found over occipital areas, theta-band activity (4–8 Hz) over frontal regions, and beta (13–30 Hz) and mu rhythms (characterised by spectral peaks in the alpha and beta ranges) around perirolandic areas, whereas delta (0.5–4 Hz) and gamma-band oscillations (>30 Hz) are found to be less conspicuous and not always localized to the same regions (Chen et al., 2008; Congedo et al., 2010; Groppe et al., 2013; Hillebrand et al., 2012; Mahjoory et al., 2020; Niso et al., 2016; Ramkumar et al., 2014).

However, beyond this general description, information regarding the characteristic oscillatory activity of individual brain regions is surprisingly scarce. Two approaches have been employed in order to achieve this aim. First, typical rhythmic patterns at specific brain locations have recently been inferred from both intracranial EEG (iEEG; Frauscher et al., 2018; Kalamangalam et al., 2020) and MEG recordings (Keitel & Gross, 2016; Mellem et al., 2017). These studies have provided the first detailed maps of ongoing rhythmic activity throughout the brain. However, a common methodological constraint is the use of predefined frequency bands and/or regions of interest. While it is a common practice to classify oscillatory activity into bands, this is not optimal since the borders between canonical frequency ranges have been arbitrarily drawn and, consequently, oscillations arising from the same physiological machinery in different species, at different ages, or even in the same individual across states are often labelled as different rhythms (Boersma et al., 2011; Buzsáki, 2006; Buzsáki et al., 2013). Similarly, in most studies the brain is parcellated into predefined regions of interest based on an anatomical atlas, which does not necessarily have to overlap with the organisation of the brain based on its functional properties (Eickhoff et al., 2018).

An alternative approach to mapping region-specific spectral patterns is based on direct perturbation of intrinsic oscillations. The stimulation of a specific cortical location by a single pulse of transcranial magnetic stimulation (TMS) evokes an oscillatory response that can be detected by concurrent EEG recordings (Ferrarelli et al., 2012; Rosanova et al., 2009). The main frequency of the evoked response is referred to as the ‘natural’ frequency and indicates the preferred rate at which the targeted cortical region is intrinsically tuned to oscillate (Basar, 1992.; Rosanova et al., 2009). These studies have accurately detected natural frequencies within the alpha range in the occipital cortex, low beta-band responses in parietal and perirolandic areas, and high beta-band oscillations in the prefrontal cortex (Ferrarelli et al., 2012; Rosanova et al., 2009). Since TMS stimulation is restricted to few locations, Amengual et al. (2019) have recently tried to develop a finer-grained atlas of natural frequencies by applying single pulses of intracranial electrical stimulation while recording iEEG in epileptic patients. Although this is a promising tool, the positioning of electrodes is constrained by clinical criteria for ethical reasons, resulting in some regions being over-sampled (e.g., temporal lobe), while others are under-sampled.

The present study aimed to address the limitations of previous research to create a detailed normative atlas of the natural frequencies of the resting human brain. Natural frequency in this context could be defined as the peak frequency of the most characteristic spectral pattern of a brain region in comparison with others. To further characterise the brain’s intrinsic rhythmic activity, we additionally identified the ‘dominant’ frequencies, i.e., the predominant oscillatory frequency of a brain region. A non-invasive strategy was employed to detect distinctive spectral patterns in a large sample of healthy individuals of a previously recorded MEG dataset. To avoid an *a priori* definition of frequency ranges and regions of interest, we applied a multivariate data-driven approach whereby exact natural frequencies were identified on a voxel-by-voxel basis.

## Methods

### Participants and data acquisition

Data were downloaded from the OMEGA database (Niso et al., 2016), which provides open access to anonymized resting state MEG data and anatomical T1-weighted Magnetic Resonance Images (MRIs). Brain activity was acquired with a whole-head CTF MEG system at the Montreal Neurological Institute (MNI, McGill University). MEG data were recorded in a magnetically shielded room with 275 axial gradiometers and 26 MEG reference sensors at a 2400 Hz sampling rate. An anti-aliasing low-pass filter at 600 Hz and CTF third-order gradient compensation were applied online. In addition, individual head shapes and three anatomical landmarks (nasion, left- and right- pre-auricular points) were digitized to facilitate MEG-MRI co-registration. Data collection and analysis were approved by the institutional ethics committees and conducted in compliance with the declaration of Helsinki.

MEG recordings were collected from participants while sitting upright. They were asked to stay awake and keep their eyes open on a fixation cross for 5 minutes. We analysed data from 128 healthy volunteers (68 males, 118 right-handed, mean age 30.5 ± 12.4 [M ± SD] years, age range 19-73 years). We only used data from one recording session per participant.

### Pre-processing

Data analysis was conducted with FieldTrip (version 20180405; Oostenveld et al., 2011) and in-house Matlab code. All scripts necessary to reproduce the analysis and the figures in this paper are available at https://github.com/necog-UAM/OMEGA-NaturalFrequencies. MEG signals were first denoised by applying Principal Component Analysis (PCA) on the reference sensors. Then, we applied a third-order Butterworth high-pass filter with a cut-off frequency of 0.05 Hz to remove slow fluctuations. Power line noise at 60 Hz, and harmonics at 120 and 180 Hz were reduced via spectrum interpolation (Leske & Dalal, 2019). Finally, the MEG signal was demeaned, detrended, and resampled at 512 Hz.

Artifact correction was performed using Independent Component Analysis (ICA), following a PCA-based dimensionality reduction to 40 components. ICs reflecting cardiac, ocular, or muscular activity were identified and projected out of the MEG signal. The data were visually inspected for any remaining artifacts. Contaminated data segments were excluded from further analysis.

### Reconstruction of source-level activity

The T1-weighted MRI of each individual was co-registered to the MEG head coordinate space by means of a semi-automatic procedure. An initial alignment was obtained by manually locating three anatomical landmarks (nasion, left- and right- pre-auricular points) in the MRI. We then employed a modified version of the Iterative Closest Point algorithm (Besl & McKay, 1992) to automatically fit the digitized head shape onto the scalp surface extracted from the participant’s MRI.

The forward model was computed using a realistic single shell volume conductor model (Nolte, 2003). We adapted a MNI-standard grid of 1-cm resolution to each individual’s brain volume and compute the lead fields for every voxel. Only voxels located within the cerebral cortex and hippocampus were considered.

Source-level time series were then reconstructed using linearly constrained minimum variance beamforming (LCMV; Van Veen et al., 1997). We employed the covariance of the artifact-free data to compute the spatial filter weights with the regularization parameter lambda set to 10%. Beamforming weights were subsequently used to estimate source-space time series from sensor-level data (Figure 1A).

**Figure 1.**
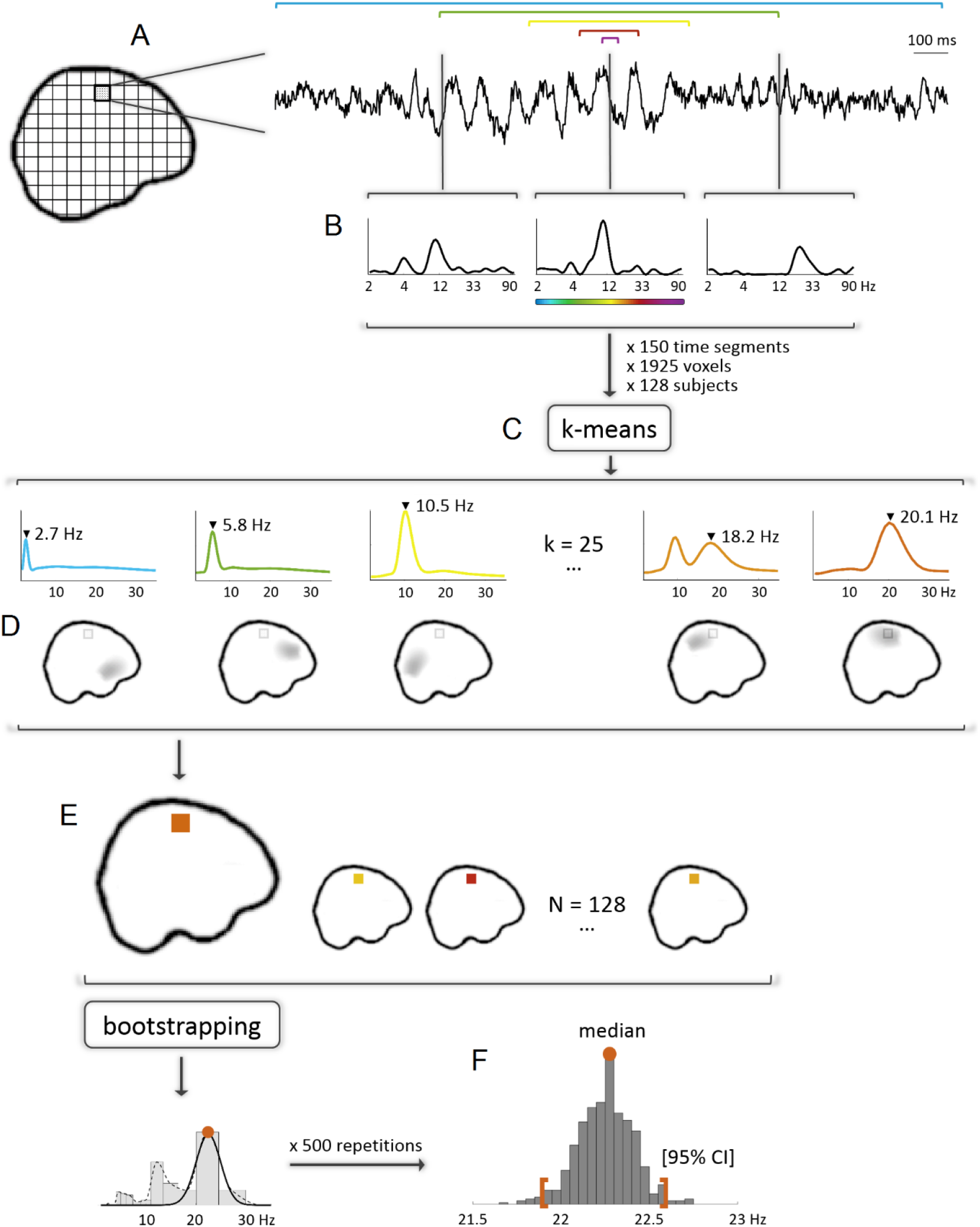
Methodological pipeline for obtaining the brain atlas of natural frequencies. **(A)** Beamforming reconstruction of source-level time series. **(B)** Spectral analysis of source-level data using a frequency-dependent window width of 5 cycles per frequency. **(C)** K-means clustering (k = 25) of source-space power spectra computed for 150 time segments x 1925 voxels x 128 subjects. **(D)** Calculation of the proportion of spectra classified within each cluster for every voxel and participant (z-scored across voxels). By averaging the obtained z-values across subjects, we identified the brain generators of each spectral pattern. **(E)** Identification of natural frequencies, defined as the peak frequency of the centroid power spectrum with the largest z-value. **(F)** Computation of the median and 95% confidence interval for each voxel’s natural frequency by means of bootstrapping.

### Frequency analysis on source-level data

Frequency analysis was conducted on the reconstructed source-space signal. We applied a Hanning-tapered sliding window Fourier transform moving in steps of 500 ms. Oscillatory power was computed for 82 frequency bins logarithmically spaced from 1.7 to 99.5 Hz. We used a frequency-dependent window width set to 5 cycles per time window, which attenuated the 1/f aperiodic component of the signal and highlighted periodic oscillatory activity. To avoid edge effects, we discarded 5.8-s time intervals (i.e., 10 cycles of the lowest frequency) at the beginning and end of every artifact-free data segment. As a result, we obtained a set of 505 ± 59 power spectra per voxel and participant, each containing the frequency components of the signal at a given time instant. Finally, in order to account for the center of the head bias, each spectrum was expressed as relative power by dividing the power at each frequency by the absolute power summed over the whole spectrum (Figure 1B).

### Cluster analysis of power spectra

We applied k-means cluster analysis to identify different patterns of source-reconstructed oscillatory activity. To lighten computational load, 150 spectra per participant were randomly selected. Then, we concatenated the power spectra of every individual (128), voxel (1925), and time segment (150) and performed k-means clustering. The cosine was employed as the distance metric to maximize differences between clusters based on the shape of the spectra (Keitel & Gross, 2016). To optimize the results, we run 5 clustering replicates with a maximum number of 200 iterations each and accepted the one with the lowest sum of distances. The number of clusters was set a priori to 25. Different choices yielded similar results, either splitting or merging adjacent clusters. We thus detected 25 different spectral patterns, most of which were characterised by a peak at a single frequency (Figure 1C).

### Brain regions underlying spectral patterns

The next step was to identify which brain regions were generating each spectral profile. For each subject and voxel, we calculated how many spectra — out of a maximum of 150 — were grouped within each cluster. To minimise the impact of spatial leakage arising from dominant sources and frequency bands (e.g., parieto-occipital alpha), values were normalised to z-scores by subtracting the mean and dividing by the standard deviation across voxels. A larger z-value at a given voxel would thus indicate that oscillatory activity in this voxel is characterised by a specific spectral pattern or, in other words, that oscillations at a specific rate are more common in this voxel than in others.

The brain generators underlying each centroid power spectrum were identified by averaging the z-values obtained for each cluster and voxel across participants (Figure 1D). Clusters reflecting gamma-band activity showed maximal z-values over the outer boundary of the cerebral cortex, surrounding the frontal, temporal, and occipital cortices. Since this result is most likely due to non-neural activity, such as microsaccades and/or electromyographic activity of facial and neck muscles (Muthukumaraswamy, 2013; Whitham et al., 2007), we only took into account clusters characterised by power spectra peaking below 30 Hz (i.e., 19 out of 25 clusters).

A non-parametric permutation test was applied to evaluate which voxels were significantly associated with every spectral pattern. For each cluster, we conducted a one-sample t-test to compare each voxel’s z-value with the mean z-value across voxels. We then estimated the distribution of test statistics under the null hypothesis by randomly shuffling (1000 times) z-values across voxels and computing t-tests against the mean z-value. In each iteration we stored the maximum t-statistic to correct for multiple comparisons. We then computed the 95th percentile of the permutation distribution. Voxels with z-values above this threshold were considered statistically significant.

### Brain map of natural frequencies

Based on the above results, we created a brain map of natural frequencies. For each voxel and participant, we identified the spectral pattern with the highest z-value. The natural frequency of a given voxel was then defined as the peak frequency of the most characteristic power spectrum (Figure 1E). Although most spectra showed one single frequency peak, in a few cases the spectrum had more than one peak, usually a combination of alpha and beta-band activity (i.e., mu rhythm). In this situation, the frequency of the beta component was determined as the characteristic frequency of the voxel, as this was the prevailing frequency over the surrounding brain areas. In addition, in the event that an isolated voxel was equally associated with two spectral patterns, its natural frequency was interpolated by taking the mode of the neighbouring voxels.

We then employed a bootstrap approach to compute the natural frequency for each voxel across the whole sample along with the 95% confidence interval. First, participants were sampled with replacement 500 times. In each repetition, we computed the histogram of natural frequencies for each voxel. We then performed a Gaussian fit of all candidates’ peak frequencies and selected the one with the highest goodness of fit. This step was computationally expensive but necessary, as simpler approaches such as computing the mean or the median across participants tend to provide solutions in the middle frequency range. The distribution of frequencies across bootstrap repetitions was used to derive the natural frequency (median) and the 95% confidence interval (2.5 and 97.5 percentiles) of each voxel (Figure 1F).

Additionally, we estimated the natural frequencies of the brain areas defined by the Automated Anatomical Labeling (AAL) atlas (Tzourio-Mazoyer et al., 2002). Although our analysis has been conducted on a single-voxel basis, identifying the typical frequency of commonly used regions of interest will enable comparison with previous studies. Thus, we detected the set of voxels located within each cortical and hippocampal region of the AAL template. Whilst the easiest approach would have then been to take the mean frequency across voxels, this was not the most appropriate strategy considering that the map of typical frequencies contains several sharp transitions that do not overlap with the AAL parcellation (e.g., theta and high-beta oscillations in the superior frontal gyrus). To overcome this difficulty, we opted for detecting a representative voxel from each AAL region by performing k-means clustering for 2 clusters on the natural frequencies of each set of voxels. We then detected which cluster contained a higher number of voxels and selected the voxel closer to the centroid as the most representative. The natural frequency and confidence interval of such voxel was then assigned to its corresponding AAL region.

### Replicability analysis

We then evaluated whether the obtained distribution of natural frequencies was replicable by repeating the above procedure in two independent subsamples of participants. We randomly split the sample into two groups (N=64) and repeat the whole analysis pipeline for each sample separately. To quantify the replicability of our results we computed the correlation coefficient between the brain maps of natural frequencies obtained for each replicate sample.

### Characterisation of spectral modes

In the preceding steps we developed a brain map of typical oscillatory activity during resting state. Thus, for every voxel in the brain we obtained a single value reflecting its natural frequency. However, ongoing oscillatory activity dynamically switches between different spectral modes (Keitel & Gross, 2016). Consequently, to provide a more comprehensive characterisation of the oscillatory profile of each brain region, we computed the median z-value for the representative voxel of each AAL region and for every canonical frequency band (delta, theta, alpha, mu, low-beta, high-beta).

Finally, we also identified the dominant frequencies of individual brain regions, i.e., the frequency that predominates most of the time in the recorded signal. For this purpose, we computed the percentage of time that oscillatory activity at every AAL brain region was clustered within each frequency band.

## Results

### Brain regions underlying spectral patterns

K-means clustering revealed 25 different spectral profiles, most of them characterised by a single peak in the power spectrum. By averaging the z-values of each cluster and voxel across participants, we identified the brain generators of each spectral pattern. As previously mentioned, we only considered clusters with peak frequencies below 30 Hz (i.e., 19 out of 25 clusters), since the generators of gamma-band oscillations were located outside the brain, surrounding frontal, temporal, and occipital regions (see Supplementary Figure 1), which is suggestive of artifactual activity.

The generators of the remaining clusters are shown in Figure 2. Only locations with z-values significantly different from zero are displayed (p < .05 corrected for multiple comparisons). As can be observed, our results show that each oscillatory frequency is generated by well-defined brain nodes. It is worth noting that although the analysis approach was purely data-driven, the frequencies associated with each brain node were naturally organized in a manner reminiscent of the canonical frequency bands: 1.5-4 Hz (delta), 4.5-7 Hz (theta), 8-13 Hz (alpha), and 14-30 Hz (beta). Not surprisingly, the dominant oscillatory frequency during the resting state was alpha. Six out of 19 clusters (31.6%) showed spectral peaks in the alpha band, in addition to 2 mu-rhythm clusters with peaks in both alpha and beta bands (10.5%).

**Figure 2.**
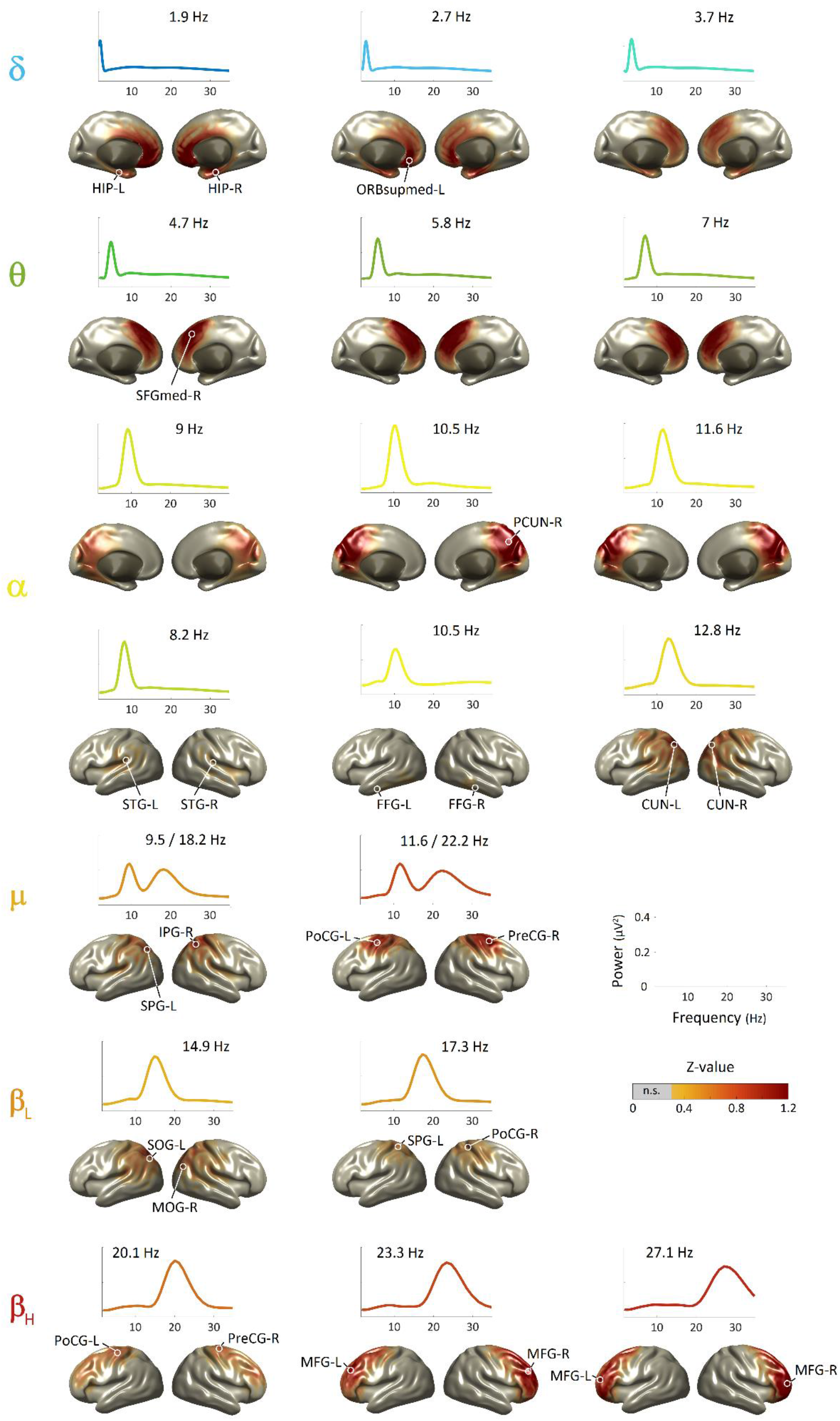
Brain generators of centroid power spectra. The figure displays the brain sources underlying the oscillatory frequencies identified by k-means clustering. Slow frequencies (i.e., delta, theta) were generated by medial fronto-temporal regions. The brain sources of alpha oscillations were twofold: a generator located at the precuneus and a set of sources distributed throughout auditory and visual cortices. The mu-rhythm was originated from somatomotor regions. Beta sources were located throughout lateral parietal and frontal cortex, following a posterior-to-anterior gradient of increasing frequency. All brain maps have been thresholded at a p-level < .05 (corrected). White circles indicate local maxima. Abbreviations: L: left; R: right; HIP: hippocampus; ORBsupmed: superior frontal gyrus (medial orbital); SFGmed: superior frontal gyrus (medial); PCUN: precuneus; STG: superior temporal gyrus; FFG: fusiform gyrus; CUN: cuneus; SPG: superior parietal gyrus; IPG: inferior parietal gyrus; PoCG: postcentral gyrus; PreCG: precentral gyrus; SOG: superior occipital gyrus; MOG: middle occipital gyrus; MFG: middle frontal gyrus.

Slow frequencies within the delta and theta range were generated by medial fronto-temporal cortical regions. In particular, we obtained three clusters with maximal z-values over the orbital part of the medial prefrontal cortex, as well as the left and right hippocampus. The corresponding power spectra peaked between 1.9 and 3.7 Hz, similar to the commonly established limits of the delta band. Three additional clusters showed spectra peaking in the theta range (4.7-7 Hz) and brain generators located around the medial superior frontal gyrus.

Interestingly, we identified two distinct varieties of alpha-band generators. First, we obtained three clusters with frequencies ranging between 9 and 11.6 Hz and a common prominent source located at the precuneus. Second, our results show a set of sources distributed bilaterally throughout sensory cortices, each of them characterised by a specific frequency peak: we found a generator peaking at 8.2 Hz over the auditory cortex, another source at 10.5 Hz around ventral visual areas, and finally a source at 12.8 Hz over dorsal visual areas.

In addition, we found two clusters with the characteristic spectrum of the mu-rhythm, i.e., a double peak in both the alpha and the beta-band (9.5/18.2 Hz and 11.6/22.2 Hz). Both sources were located over somatomotor cortex.

Finally, the brain sources of beta oscillations were distributed throughout lateral parietal and frontal regions, with increasing frequency following a posterior-to-anterior gradient. Specifically, low beta-band activity (14.9-17.3 Hz) was mostly generated by lateral occipito-parietal regions, whereas high beta-band oscillations were detected over motor (20.1 Hz) and prefrontal cortices (23.3-27.1 Hz).

### Brain map of natural frequencies

The map of natural frequencies revealed a region-specific distribution of oscillatory activity characterised by both a medial-to-lateral and a posterior-to-anterior gradient of increasing frequency. As can be observed in Figure 3A, medial frontal and temporal regions were characterised by lower rhythms (delta- and theta-band), posterior occipito-temporal cortices were dominated by alpha-band activity, and parietal areas mostly exhibited low beta-band activity. In contrast, motor and lateral prefrontal cortex were distinguished by high beta activity. The typical frequency of each voxel usually remained within a given frequency band as revealed by the 95% confidence interval maps (Figure 3B).

**Figure 3.**
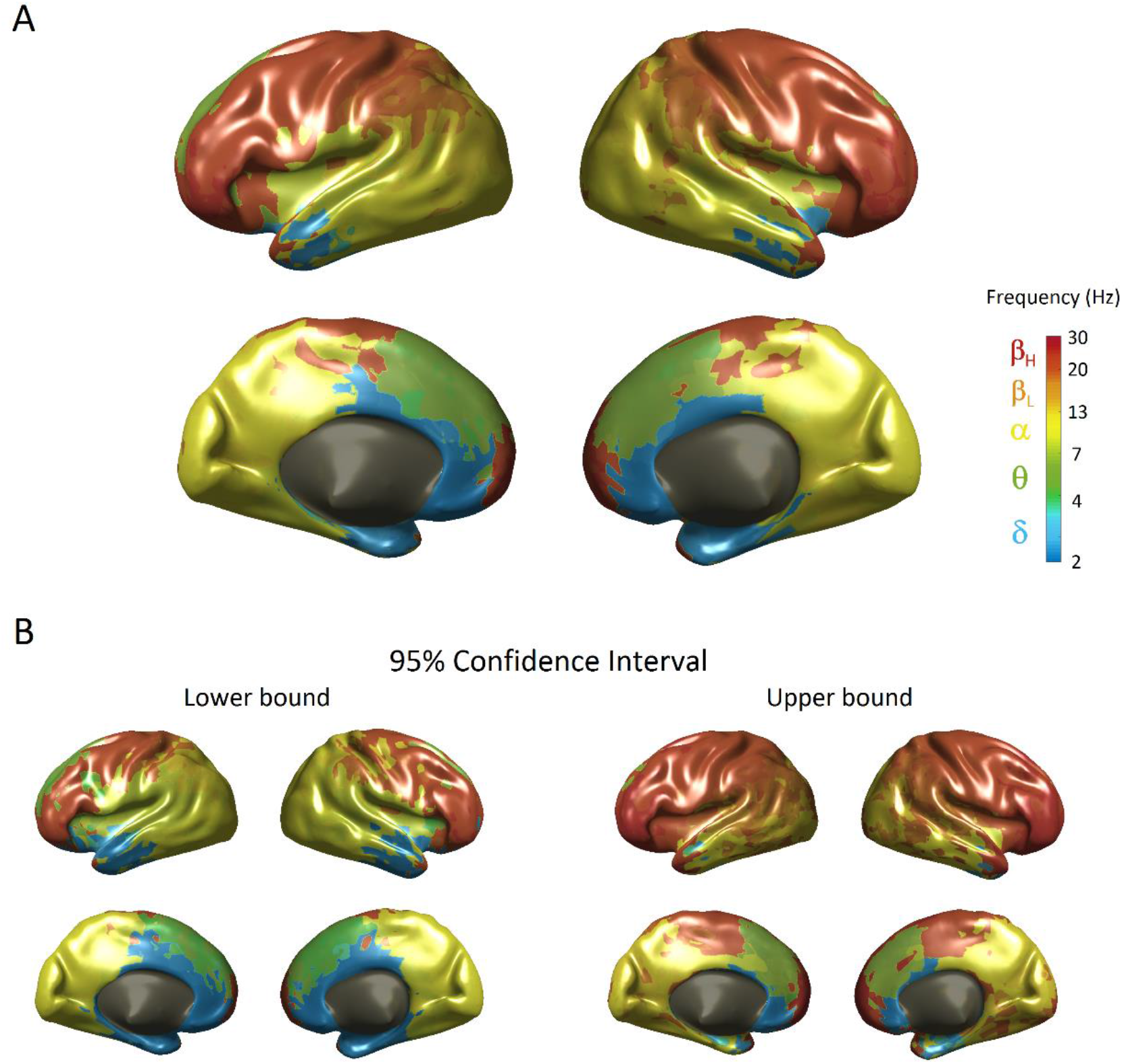
Brain map of natural frequencies during resting. **(A)** Distribution of natural oscillations at the single-voxel level exhibiting both a medial-to-lateral and a posterior-to-anterior gradient of increasing frequency. Canonical frequency band ranges are color-coded to facilitate interpretation. **(B)** Lower and upper bounds of the 95% confidence interval estimated by a bootstrapping approach.

In addition to the cortical surface representation, we have interpolated the results in a 3D volume in MNI space, creating an atlas that allows to look up the natural frequency, as well as the lower and upper bounds of the 95% confidence interval of specific MNI coordinates. The NIfTI files necessary to visualize the atlas can be found in the Supplementary Material. We have also created a Matlab function to easily retrieve the natural frequency and confidence interval at a given MNI coordinate (NaturalFreq.m, see Supplementary File 1).

Finally, the natural frequency and corresponding confidence interval for the cortical and hippocampal regions defined by the AAL atlas are listed in Table 1.

**Table 1.** Natural frequency of AAL regions. Median natural frequency and 95% confidence interval of cortical and hippocampal AAL regions.

| **AAL region** | **Natural Frequency** | **95% Confidence Interval** |
| --- | --- | --- |
| Precentral gyrus | 22.3 | [22.0, 22.6] |
| Superior frontal gyrus (dorsolat) | 23.1 | [22.7, 23.7] |
| Superior frontal gyrus (orb) | 23.3 | [22.7, 25.5] |
| Middle frontal gyrus | 23.2 | [22.7, 23.7] |
| Middle frontal gyrus (orb) | 25.5 | [23.7, 26.1] |
| Inferior frontal gyrus (oper) | 22.6 | [22.2, 23.1] |
| Inferior frontal gyrus (tri) | 23.4 | [23.0, 23.9] |
| Inferior frontal gyrus (orb) | 23.2 | [22.7, 25.3] |
| Rolandic operculum | 8.4 | [8.3, 18.1] |
| Supplementary motor area | 5.5 | [5.3, 5.7] |
| Olfactory cortex | 2.2 | [2.1, 2.3] |
| Superior frontal gyrus (med) | 6.6 | [5.6, 6.8] |
| Superior frontal gyrus (med orb) | 23.4 | [22.9, 25.5] |
| Rectus gyrus | 2.1 | [2.1, 2.2] |
| Insula | 23.0 | [22.3, 25.6] |
| Anterior cingulate gyrus | 5.5 | [4.6, 6.3] |
| Middle cingulate gyrus | 8.5 | [8.3, 8.8] |
| Posterior cingulate gyrus | 10.2 | [9.9, 10.5] |
| Hippocampus | 2.4 | [2.3, 2.5] |
| Parahippocampal gyrus | 2.3 | [2.1, 2.4] |
| Calcarine cortex | 11.5 | [10.3, 11.9] |
| Cuneus | 10.5 | [10.2, 11.7] |
| Lingual gyrus | 10.2 | [8.8, 11.7] |
| Superior occipital gyrus | 11.4 | [10.3, 11.9] |
| Middle occipital gyrus | 11.9 | [10.4, 12.3] |
| Inferior occipital gyrus | 10.4 | [10.2, 11.6] |
| Fusiform gyrus | 2.3 | [2.2, 2.4] |
| Postcentral gyrus | 21.4 | [20.8, 21.9] |
| Superior parietal gyrus | 18.7 | [18.2, 19.2] |
| Inferior parietal gyrus | 21.3 | [19.0, 22.0] |
| Supramarginal gyrus | 19.1 | [18.5, 21.1] |
| Angular gyrus | 12.1 | [11.8, 12.4] |
| Precuneus | 10.6 | [10.4, 11.6] |
| Paracentral lobule | 21.5 | [19.1, 22.2] |
| Heschl gyrus | 8.4 | [8.3, 9.6] |
| Superior temporal gyrus | 8.7 | [8.4, 9.9] |
| Temporal pole (sup) | 22.7 | [22.1, 23.3] |
| Middle temporal gyrus | 10.2 | [10.1, 10.4] |
| Temporal pole (middle) | 22.6 | [21.9, 23.4] |
| Inferior temporal gyrus | 10.1 | [9.9, 10.3] |

### Replicability analysis

The correlation coefficient between the brain maps of natural frequencies derived from two independent subsamples (N = 64) was 0.81 (p < .001), indicating that the replicability of our results is considerably high (see Figure 4).

**Figure 4.**
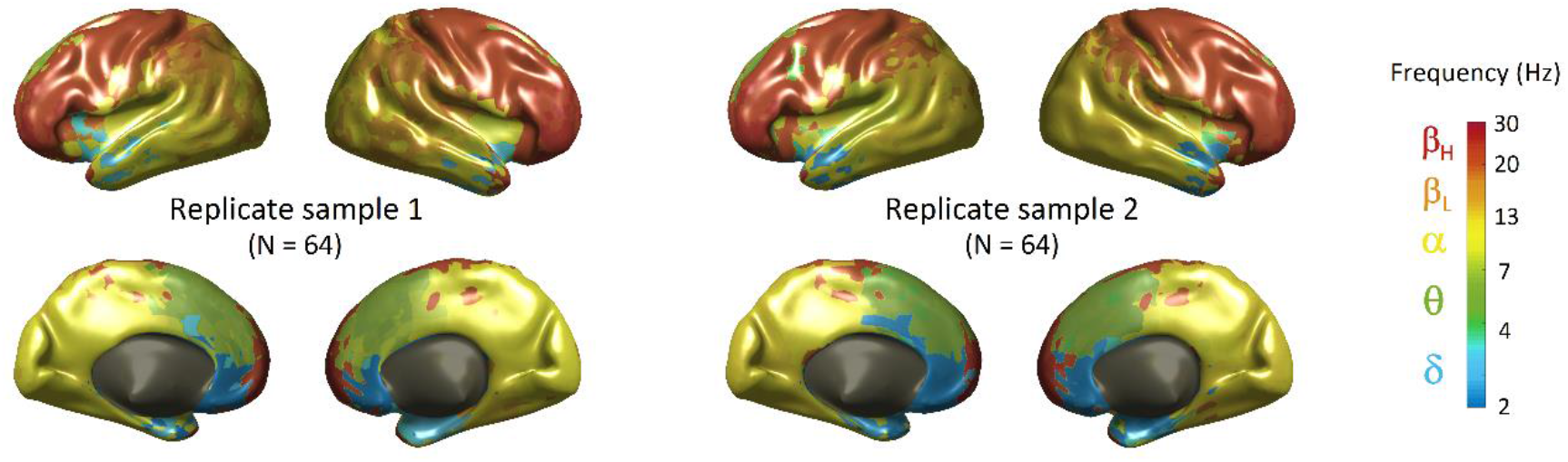
Replicability of the brain map of natural frequencies. The figure shows the brain maps estimated from two independent subsamples. The correlation coefficient between them was 0.81.

### Characterisation of spectral modes

Finally, we computed the weights of different spectral modes to further characterise the oscillatory profile of each brain area beyond its natural frequency. As can be observed in Figure 5A, most AAL regions showed more than one characteristic spectral mode. In some cases, these reflected contiguous frequency bands (e.g., delta and theta in the rectus gyrus, or alpha/mu/low-beta in the superior parietal gyrus). In other cases, different spectral modes might reflect the natural frequencies of several functional sub-parcellations of one single region (e.g., theta and high-beta along the superior frontal gyrus), as the anatomical boundaries of the AAL atlas did not completely fit with the transitions between natural frequencies. It is also worth noting that some regions, particularly throughout the temporal lobe (e.g., middle and inferior temporal gyrus), did not exhibit typical oscillatory frequencies at any of the canonical frequency bands.

**Figure 5.**
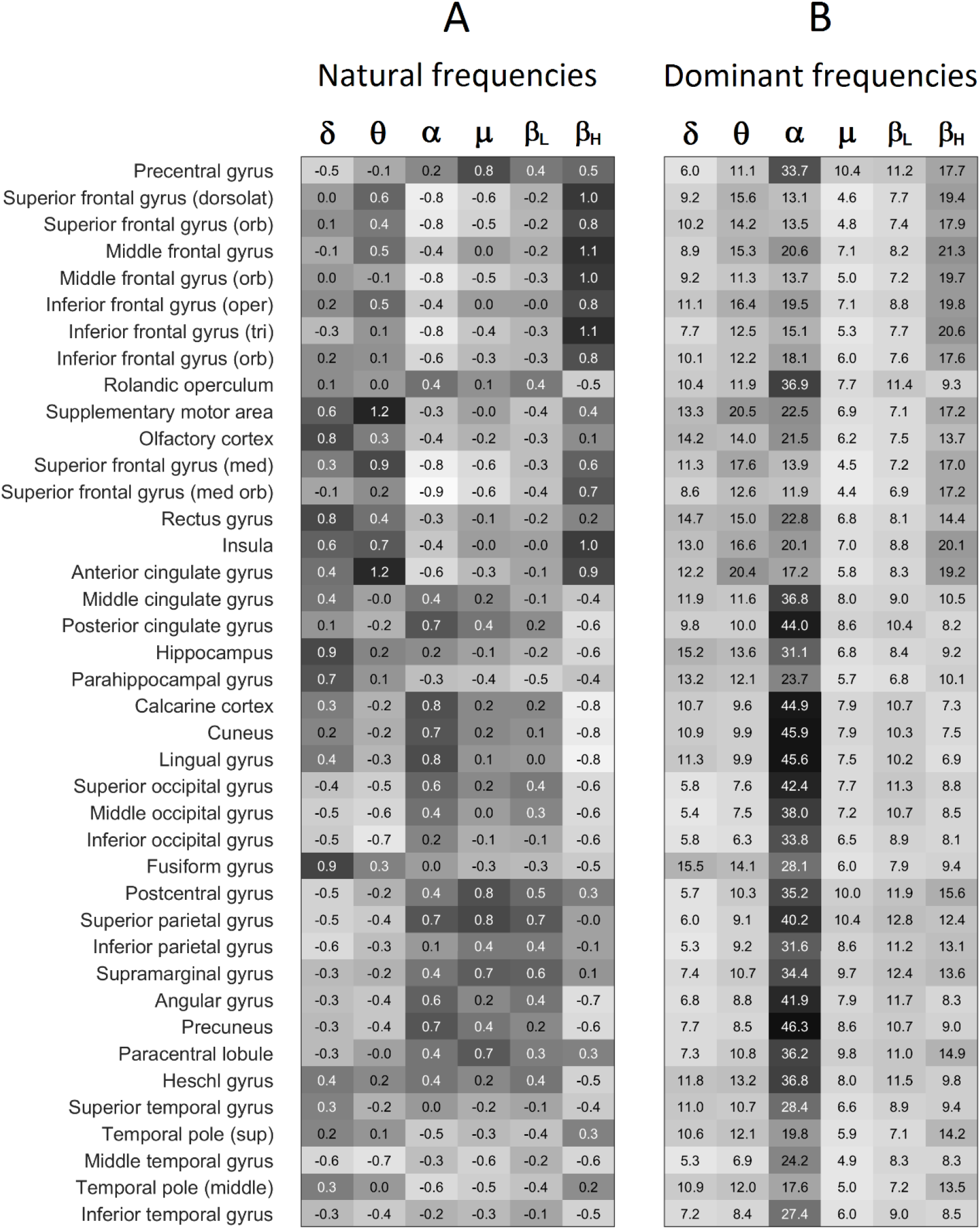
Spectral modes for natural and dominant frequencies at different AAL brain regions. **(A)** Natural frequencies were indexed by z-values (i.e., how typical is activity in a given frequency band at a specific region compared with other regions). Note that most AAL brain areas were engaged in several characteristic spectral modes. **(B)** Dominant frequencies were computed as the percentage of time spent in each spectral mode (i.e., how predominant is activity in a given frequency band). The figure shows the high predominance of alpha-band activity, particularly in posterior regions.

To complement the characterisation of ongoing oscillatory activity, we additionally identified the dominant frequencies of each AAL brain region by computing the percentage of time spent in each spectral mode (Figure 5B). Contrasting with the heterogeneity observed for natural frequencies (Figure 5A), brain activity was mostly dominated by a single mode in the alpha frequency range, especially in the posterior cortex. Dominant frequencies in frontal areas were found to be more evenly distributed among theta, alpha and beta bands.

## Discussion

This study presents the most comprehensive map of the typical oscillatory activity of the resting human brain performed to date. We applied a fully data-driven procedure to identify precise natural frequencies at a single-voxel level. Our results reveal a region-specific organisation of ongoing oscillations defined by both a medial-to-lateral and a posterior-to-anterior gradient of increasing frequency. In particular, medial fronto-temporal areas were characterised by slow (delta/theta) rhythms. Posterior regions exhibited natural frequencies within the alpha band, with specific generators in the precuneus, visual, and auditory sensory cortices. Somatomotor regions showed characteristic mu and low beta activity, while the lateral prefrontal cortex was distinguished by oscillations in the high beta range. Critically, the brain atlas of natural frequencies proved to be highly reproducible in two independent subsamples of individuals.

In the following paragraphs, we will first discuss the critical methodological features of the present study in comparison with prior work. We will then address the distinction between the concepts of natural frequency and dominant frequency. Subsequently, we will examine the results obtained for each frequency band. Whereas some results corroborate well established knowledge, such as the neural underpinnings of alpha, mu, and theta rhythms, others provide novel information on frequency bands on which there is less consensus, such as delta or beta. Finally, we will discuss the potential relationship between natural frequencies and resting state networks (RSNs).

### Methodological remarks

The present study combined a series of methodological innovations that, taken together, significantly contribute to the characterisation of intrinsic brain oscillations. First, analyses were performed on MEG data obtained from a very large (N=128) and representative (age range 19-73 years) sample of healthy adult volunteers. The recently developed iEEG atlas of electrophysiological brain activity has also been created from a large database, in this case recorded from patients with epilepsy (Frauscher et al., 2018; Kalamangalam et al., 2020). However, the ethical constraints associated with electrode placement make it more difficult to systematically map oscillatory activity throughout the brain by means of iEEG.

Second, according to our data-driven strategy, and in contrast to previous work (Keitel & Gross, 2016; Mahjoory et al., 2020), we did not parcellate the brain into predefined regions of interest based on anatomical criteria. To the best of our knowledge, this is the first study to characterise typical oscillatory activity on a voxel-by-voxel basis. Our results reveal that the brain displays similar rhythmic patterns in neighbouring voxels, which could be grouped into functionally equivalent areas (see atlas NaturalFreq.nii in Supplementary File 1). Importantly, the anatomical and functional parcellations do not completely overlap, as some abrupt transitions in natural frequencies do not have an anatomical correspondence. Future research could extend the current results to elaborate a functional parcellation based on the brain’s intrinsic oscillatory dynamics, similar to the functional organisation of the brain derived from fMRI resting state connectivity analysis over the last decade (Eickhoff et al., 2018; Yeo et al., 2011).

Finally, unlike most previous work, we did not employ predetermined frequency bands. Instead, we identified exact frequency peaks in power spectra centroids obtained from k-means clustering. Critically, spectral peak detection has recently been emphasized as a requisite to verify the presence of true oscillatory activity, as the average power computed over a narrowband frequency range does not necessarily reflect the existence of an underlying oscillation (Donoghue et al., 2021). Nevertheless, despite not having defined frequency bands *a priori*, they emerged empirically from the data. Based on their underlying brain generators, centroid power spectra could be organized into the following frequency ranges: 1.5-4 Hz for delta, 4.5-7 Hz for theta, 8-13 Hz for alpha, and 14-30 Hz for beta. These fit well with the standard classification of brain rhythms as well as with the empirical definition of frequency bands obtained with other methodological approaches (Mahjoory et al., 2020).

### Natural frequencies Vs. Dominant frequencies

A fundamental distinction must be made between the concepts of dominant and natural frequency. On the one hand, dominant frequency could be defined as the most prevalent oscillatory rhythm of a brain region, i.e., the frequency that predominates most of the time. On the other hand, natural frequency would refer to the spectral pattern that characterises an individual brain area or, in other words, the oscillatory frequency that is most often observed in a given region in comparison with other parts of the brain. As our results show, dominant and natural frequencies do not necessarily coincide (see Figure 5). An example is the tau rhythm (~8.5 Hz) which albeit not dominant, is the characteristic local rhythm of the auditory cortex as corroborated by its reactivity to sounds in previous research (Lehtelä et al., 1997; Tiihonen et al., 1991).

At a methodological level, the common approach of averaging power spectra tends to emphasise dominant frequencies (Hillebrand et al., 2012; Mahjoory et al., 2020; Mellem et al., 2017; Niso et al., 2016). In contrast, clustering-based techniques allow the identification of non-dominant patterns present on single trials, thus enabling the identification of natural frequencies (Frauscher et al., 2018; Keitel & Gross, 2016). However, a common problem is that a highly dominant source, such as the posterior alpha rhythm, may obscure oscillatory activity in neighbouring regions. In this study, we therefore applied z-score normalisation to mitigate the potential confounding effect of source leakage.

The concept of natural frequency has not extensively been used in the literature. Until now, this concept has almost exclusively been employed to refer to the main frequency evoked by direct electrical/magnetic stimulation of a given brain region (Amengual et al., 2019; Ferrarelli et al., 2012; Rosanova et al., 2009). In this study, we have developed a map of natural frequencies that is highly compatible with the results obtained by direct cortical perturbation, but without interfering with ongoing brain activity. For example, our results are very similar to those obtained by Rosanova and colleagues (2009), who found natural frequencies for the occipital cortex at 10.8 Hz (11.4 Hz for an equivalent region in our study), for the superior parietal cortex at 18.6 Hz (18.7 Hz here), and for the premotor cortex at 29 Hz (23.1 Hz here). Our results are also broadly consistent with the natural frequencies reported by Ferrarelli et al. (2012) in the parietal cortex (19.9 Hz Vs. 18.7 Hz here), the motor cortex (21.8 Hz Vs. 22.3 Hz here), the premotor cortex (26.5 Hz Vs. 23.1 Hz here), and the prefrontal cortex (31 Hz Vs 23.2 Hz here). It is worth noting that an advantage of our approach is that it allows the extraction of whole-brain natural frequencies with a wide and homogeneous spatial coverage from 5-minute resting state recordings.

### Brain generators underlying canonical frequency bands

#### Default-alpha Vs. sensory-specific alpha oscillations

Unsurprisingly, our first finding was that the alpha rhythm is clearly dominant in the brain: most clusters were characterised by an alpha peak (see Figure 2) and for most of the time rhythmic brain activity oscillated around 10 Hz in the majority of brain regions, with the exception of frontal areas (Figure 5B). Alpha oscillations have traditionally been divided into two types, posterior-alpha and central-alpha (i.e., mu rhythm). However, our map of natural frequencies shows a much greater richness of oscillators in the alpha range, in line with current views (Hindriks et al., 2017; Ramkumar et al., 2014).

A first type of alpha generator, peaking at around 10.5 Hz, was located in the precuneus, in agreement with pioneering work that identified this brain area as the main source of posterior alpha activity (Hari et al., 1997; Salmelin & Hari, 1994). Further, the precuneus/posterior cingulate cortex (PCC) is considered a key hub of the default mode network (DMN) (Fransson & Marrelec, 2008; Raichle et al., 2001), with the highest resting metabolic rate within this network (Gusnard & Raichle, 2001). Given that the relationship between alpha and the DMN has been well established by previous studies (de Pasquale et al., 2012; Knyazev et al., 2011; Mantini et al., 2007; Rusiniak et al., 2018; Scheeringa et al., 2012), this generator could be referred to as ‘default-alpha’.

We found a second type of alpha-band generators distributed over visual, auditory and somatosensory (linked to mu-rhythm) cortices, in agreement with prior evidence (Haegens et al., 2015; Hari & Salmelin, 1997; Salmelin & Hari, 1994). Each sensory-specific alpha was characterised by a particular cortical distribution and natural frequency. Thus, in the upper alpha range (10.5-13 Hz), ‘visual-alpha’ was distributed throughout the ventral and dorsal visual streams, similar to the results obtained with diverse methodological tools (Barzegaran et al., 2017; Hindriks et al., 2017). We also identified a source of alpha oscillations over the auditory cortex — also referred to as ‘tau’ rhythm — with a lower natural frequency at around 8.5 Hz, as expected on the basis of previous work (Lehtelä et al., 1997; Ramkumar et al., 2014; Tiihonen et al., 1991).

It is important to highlight that what is commonly known as posterior-alpha may be an overlay of two generators with similar topographic distribution (but functionally very different), that is, the default-alpha and the visual-alpha. An increase in default-alpha activity would be related to internal mental processing and concurrent reduced attention to external stimuli, which would consequently lead to a decrease in visual-alpha activity (Knyazev et al., 2011). This idea is supported by studies showing that alpha power tends to correlate positively with blood oxygen level-dependent (BOLD) activity in the DMN, while there is a negative correlation in visual areas (Mantini et al., 2007; Scheeringa et al., 2012). It is important to keep in mind that current analyses have been conducted on resting state data. It would be interesting for future studies to extend the present map of ongoing natural frequencies to various perceptual and cognitive tasks, to test whether or not natural frequencies are state-dependent.

#### Somatomotor mu rhythm

We found somatosensory alpha activity linked to the mu rhythm over perirolandic cortex. Mu oscillations are characterised by a comb-shaped waveform and a spectrum with a double peak in the alpha and beta bands (Tiihonen et al., 1989). Initially, MEG studies suggested that the mu rhythm arises from the superposition of a mu-alpha component originating in the somatosensory cortex and a mu-beta component reflecting activity from the precentral motor cortex (Hari & Salmelin, 1997). However, iEEG recordings have shown that both precentral and postcentral regions present modulations of alpha as well as beta oscillations (Crone et al., 1998), consistent with the observation that the mu rhythm is reactive to both movement and tactile stimulation (Kane et al., 2017). Our results are in line with the latter view, as we found a mu rhythm with both alpha and beta spectral components widely distributed over the somatomotor cortex.

Interestingly, previous research has demonstrated that mu-alpha and mu-beta components can be dissociated, with only the alpha or only the beta component appearing up to 50% of the time (Jones et al., 2009). This might explain why in our study, precentral and postcentral areas, though mainly characterised by mu oscillations, also displayed spectral modes in the alpha and beta bands (see Figure 5A). Furthermore, since we could only assign one frequency value per voxel to create the map of natural frequencies, we were compelled to select either the mu-alpha or the mu-beta frequency peak. Although mu is usually considered to belong to the family of alpha oscillations, neighbouring brain areas predominantly exhibited beta activity and, therefore, the mu-beta peak prevailed over mu-alpha. This decision was supported by previous evidence showing a predominance of beta activity over perirolandic cortex (Jasper & Penfield, 1949; Mantini et al., 2007), as well as the recent characterisation of somatosensory activity as transient beta events that may or may not appear embedded within an alpha oscillation (Sherman et al., 2016).

#### Midfrontal theta oscillations

Theta oscillations are often observed at the scalp level with a characteristic topographical distribution, centred over midline fronto-central sites (Scheeringa et al., 2008). Brain sources underlying midfrontal theta have been consistently located near dorsal anterior cingulate cortex (ACC) or medial prefrontal cortex (Gevins et al., 1997; Onton et al., 2005). This is highly consistent with the brain regions showing natural frequencies within the theta range found here, as well as with the location of the resting state theta-band generators reported by previous studies (Kalamangalam et al., 2020; Mellem et al., 2017; Niso et al., 2016). Moreover, midfrontal theta oscillations are commonly induced by highly demanding cognitive tasks, such as mental arithmetic, error detection, response conflict, or working memory tasks (Gartner et al., 2015; Gevins et al., 1997; Zuure et al., 2020). Taking into consideration both anatomical and functional evidence, it might be suggested that theta oscillations arising from medial prefrontal cortex underlie the RSN of executive control and working memory previously identified with fMRI (Cavanagh & Frank, 2014; Damoiseaux et al., 2006).

#### Orbitofrontal and hippocampal delta-band oscillations

The two brain areas characterised by natural frequencies in the delta band were the medial orbitofrontal cortex and the hippocampal region. The orbitofrontal cortex has been pointed out as generator of slow oscillations at rest (Congedo et al., 2010; Hillebrand et al., 2012; Niso et al., 2016) and during sleep, in which they propagate as travelling waves towards posterior regions periodically modifying the state of cortical excitability and connectivity (Massimini et al., 2004). Even though this generator needs to be interpreted with caution as it might be related to ocular artifacts (Ramkumar et al., 2014), delta oscillations have also been identified in orbitofrontal cortex with iEEG (Frauscher et al., 2018), that is less prone to contamination by eye-movements.

The hippocampus and parahippocampal gyrus showed natural frequencies in the delta range, around 2-3 Hz. This finding is consistent with the iEEG atlas developed by Frauscher and colleagues (2018), although it seems at first to contradict the common notion that theta oscillations underlie hippocampal function (Herweg et al., 2020). However, this apparent contradiction might be overcome by evidence that theta oscillations become slower with increasing brain size (Buzsáki et al., 2013). Thus, rodent theta oscillations (~4-10 Hz) would be expressed in humans at lower frequencies, within the delta band (~1-4 Hz) (Jacobs, 2014). Critically, hippocampal delta oscillations have been observed in intracranial recordings in humans during the performance of spatial navigation (Watrous et al., 2013a) and memory tasks (Lega et al., 2012), as well as during REM sleep (Bódizs et al., 2001), supporting the notion that human-delta and rodent-theta might be functionally equivalent (Jacobs, 2014).

It could also be possible that both delta and theta-band oscillations coexist in different parts of the hippocampus. In fact, we found a sharp transition between the anterior portion of the hippocampus, with natural frequencies within the delta band (~2-2.5 Hz), and the posterior portion, characterised by oscillations in the upper theta range (~8-8.5 Hz) (see NaturalFreq.nii in Supplementary File 1). Since the anterior and posterior hippocampus are known to be functionally distinct (Fanselow & Dong 2010; Zeidman, 2016), an intriguing possibility is that this dissociation is expressed in specific oscillatory dynamics. In support of this idea, some studies have identified two different types of hippocampal oscillations at 2-3 Hz and ~8 Hz during memory processes (Lega et al., 2012; Watrous et al., 2013b). However, given the limited spatial resolution of MEG, further investigation with iEEG would be necessary to confirm this finding.

#### Lateral prefrontal beta-band oscillations

Beta oscillations have traditionally been considered a signature of the resting motor cortex (Jasper & Penfield, 1949). Most previous research supports this view, showing that beta activity mainly arises from perirolandic regions (Frauscher et al., 2018; Hillebrand et al., 2012; Niso et al., 2016). Our results only partially agree with this notion, since we found that the beta rhythm characterises the entire lateral surface of the frontal lobe. Thus, perirolandic cortex would be distinguished by natural frequencies around 20 Hz — often in the form of mu oscillations as previously discussed —, whereas premotor and lateral prefrontal regions would exhibit natural frequencies in the upper beta range, between 20 and 30 Hz.

Only a few previous studies have also pointed to beta as the characteristic rhythm of the prefrontal cortex (Mahjoory et al., 2020; Mellem et al., 2017). This could be due to the fact that, unlike posterior regions, the prefrontal cortex does not show a clear dominant spectral mode (see Figure 5B). The different methodologies employed in the various studies may have emphasised different spectral modes, leading to disparate results regarding the typical frequency of this region.

Furthermore, oscillatory activity in the lateral prefrontal cortex is not homogeneous but rather characterised by a posterior-to-anterior gradient of increasing frequency, in agreement with prior reports (Ferrarelli et al., 2012; Mahjoory et al., 2020). Intriguingly, this gradient is reminiscent of Koechlin’s theory of cognitive control, which considers that the lateral prefrontal cortex is functionally organised as a cascade of executive processes (sensory, contextual, episodic, and branching control) that are hierarchically ordered along the anterior-posterior axis (Koechlin & Summerfield, 2007). Therefore, it might be speculated that a progressively increasing frequency within the beta band indexes the multistage functional architecture of the lateral prefrontal cortex.

### Natural frequencies and Resting State Networks (RSNs)

Given the pivotal role of brain oscillations in brain communication (Buzsáki & Watson, 2012; Fries, 2015; Varela et al., 2001), an arising question is whether natural frequencies are somehow related to network connectivity. At first, one might expect different nodes of the same RSN to share the same natural frequency in order to facilitate synchronization. However, our results do not support this idea. For example, the main nodes of the DMN are characterised by natural frequencies in virtually all frequency bands: alpha for the PCC, delta/theta for the medial prefrontal cortex, and beta for the inferior parietal gyrus. Nonetheless, a mismatch in natural frequencies does not preclude network communication, as will be discussed next.

Communication between brain areas can be attained through different mechanisms which may act at different timescales: one mechanism based on amplitude correlation for slower timescales and another one based on phase coherence for fast-resolved communication (Siegel et al., 2012). Firstly, amplitude correlation of band-limited power has been proposed as a mechanism for large-scale network communication, which is often observed within specific frequency bands, particularly in the alpha and beta ranges (de Pasquale et al., 2012; Brookes et al., 2011). It would be interesting for future studies to additionally explore potential cross-frequency amplitude correlations of power at the natural frequencies of RSN nodes, to thereby investigate whether natural frequencies play a role in this mechanism of brain communication.

Secondly, network communication could also take place via phase coherence, either at equal or at different frequencies (i.e., cross-frequency coupling). On the one hand, brain regions might transiently abandon their natural frequencies, adopting a different spectral mode to synchronise at the same frequency. This would be in line with the recent view that RSNs are non-stationary, but rather characterised by short-lived transient brain states (Vidaurre et al., 2018). During these microstates, different parts of the network may synchronize at a particular frequency. For instance, it has been found that the anterior and posterior portions of the DMN synchronise at the typical frequency of the anterior region, i.e., in the delta/theta range (1-7 Hz) (Vidaurre et al., 2018). Thus, the PCC might briefly leave its natural frequency in the alpha range to communicate with the anterior DMN in a slower frequency channel. Our results are compatible with this scenario, since all brain regions show activity in several spectral modes beyond their natural frequency (Figure 5A; see also Keitel & Gross, 2016), which is suggestive of a dynamically changing state of the oscillatory brain activity.

On the other hand, communication can also be achieved through phase-phase or phase-amplitude coupling at different frequencies (Palva & Palva, 2018; Roopun et al., 2008). For example, communication between the different nodes of the DMN characterised by natural frequencies at ~5, ~10, and ~20 Hz could take place via 1:2 and 1:4 cross-frequency coupling (Palva & Palva, 2018). Further, it has been proposed that the hierarchical organisation of oscillations at multiple timescales might constitute the cornerstone of the neural syntax (Buzsáki & Watson, 2012) and, therefore, a deeper understanding of typical intrinsic oscillatory activity could contribute to unravel the brain’s communication code.

### Conclusions

Here we have presented the first MEG-based atlas of the natural frequencies of the human brain at rest. Overall, our results show that the brain exhibits a characteristic and highly replicable pattern of ongoing oscillatory activity, which is organised according to both an anterior-posterior and a medial-lateral axis. We believe that these findings will contribute to further elucidate the functional architecture of the human brain.

## Acknowledgements

We are thankful to the team of researchers and technicians involved in The Open MEG Archive (OMEGA) project for recording and making publicly available the database employed in the present study and, particularly, to Guiomar Niso for introducing us to this initiative. The authors also thank Abel Cano for valuable advice regarding the use of the k-means algorithm. This work was supported by FEDER/Ministerio de Ciencia, Innovación y Universidades – Agencia Estatal de Investigación, Spain (grant PGC2018-100682-B-I00).

