## Supplementary Figure 1 for "The natural frequencies of the resting human brain: an MEG-based atlas"

### Supplementary Material

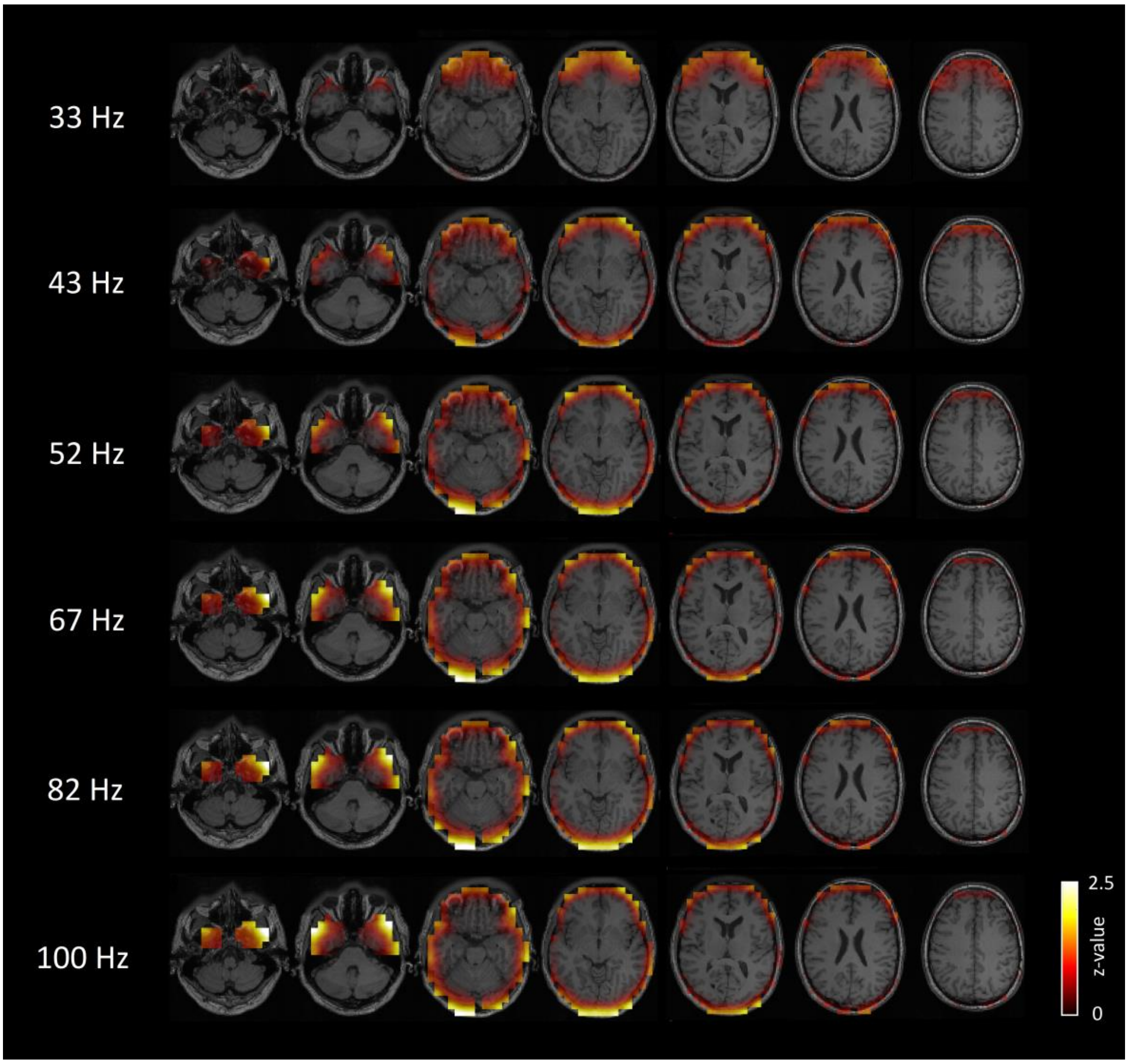


**Suppl. Figure 1. Brain generators of centroid power spectra peaking above 30 Hz.** Voxels showing characteristic activity within the gamma band were located outside the brain, surrounding the frontal, temporal, and occipital cortices. This pattern is suggestive of artifactual ocular and/or muscular activity.
