## Supplementary File 1 for "The natural frequencies of the resting human brain: an MEG-based atlas"

### Supplementary Material

**Suppl. File 1. Zip file containing the atlas of natural frequencies in NIfTI format and a Matlab function developed to retrieve natural frequencies at specific MNI coordinates.** The file NaturalFreq.nii allows visualization of the atlas of natural frequencies of the resting human brain at the voxel level. NaturalFreq_Low95%CI.nii and NaturalFreq_High95%CI.nii indicate the lower and upper bounds of the 95% confidence interval, respectively. The function NaturalFreq.m provides information of the natural frequency and the corresponding confidence interval at any given MNI coordinate. File available at https://github.com/necog-UAM/OMEGA-NaturalFrequencies.
